# Feeding schedule and proteolysis regulate autophagic clearance of mutant huntingtin

**DOI:** 10.1101/116178

**Authors:** Dagmar E Ehrnhoefer, Dale DO Martin, Xiaofan Qiu, Safia Ladha, Nicholas S Caron, Niels H Skotte, Yen TN Nguyen, Sabine Engemann, Sonia Franciosi, Michael R Hayden

## Abstract

The expression of mutant huntingtin (mHTT) causes Huntington disease (HD), and lowering its levels is therefore an attractive therapeutic strategy. Here we show that scheduled feeding significantly decreases mHTT protein levels through enhanced autophagy in the CNS of an HD mouse model, while short term fasting is sufficient to observe similar effects in peripheral tissue. Furthermore, preventing proteolysis at the caspase-6 cleavage site D586 (C6R) makes mHTT a better substrate for autophagy, additionally increasing its clearance. Mice expressing mutant C6R also exhibit increased autophagy at baseline compared to an HD model with cleavable mHTT, suggesting that the native function of HTT in promoting autophagy is disrupted upon cleavage and re-established by prevention of cleavage by caspase-6. In HD patients, mHTT clearance and autophagy may therefore become increasingly impaired as a function of age and disease stage by gradually increased activity of mHTT-processing enzymes.

## Introduction

Huntington disease (HD) is an autosomal dominant neurodegenerative disorder that is caused by an expansion of a polyglutamine tract in the huntingtin (HTT) protein (1). Mutant HTT (mHTT) causes dysfunction in different cellular compartments (2) that are difficult to target separately. The removal of mHTT itself is therefore an attractive therapeutic strategy and is currently being pursued in both clinical and pre-clinical studies (3). While most of these studies aim to lower HTT RNA, changes in mHTT protein levels through increased degradation have also been shown to ameliorate HD symptoms (4, 5). Both soluble and aggregated forms of mHTT are thought to be cleared preferentially through autophagy (6, 7), and both mTOR-dependent and -independent autophagic pathways have been implicated in its degradation (4, 8). This suggests that mHTT degradation is largely independent of the initial autophagy-inducing stimulus.

Autophagy is strongly influenced by metabolic status and circadian rhythms (9-11), both of which are altered in HD (12-16). Many pre-clinical studies using mTOR inhibitors or AMPK activators, which modulate major pathways sensing cellular energy status, have shown beneficial effects on HD phenotypes (5, 17, 18). Additionally, sleep dysfunction is a prominent symptom in human HD patients (13, 19), and therapeutic interventions to normalize circadian rhythm have been suggested, but have yet to be systematically evaluated. However, studies in the R6/2 mouse model of HD have demonstrated that scheduled feeding forces the normalization of circadian rhythms and can ameliorate motor phenotypes (12, 20).

We therefore decided to investigate the impact of fasting and scheduled feeding on autophagy pathways and mHTT clearance. We show that regulated food intake activates autophagy in the YAC128 mouse model of HD and reduces the levels of mHTT protein in the brain, providing a potential molecular mechanism for the reported beneficial effects of this intervention.

In addition, we demonstrate that the autophagic clearance of mHTT is also increased by preventing its proteolysis at amino acid 586 in the C6R mouse model, an *in vivo* system in which mHTT does not elicit any known HD-related phenotypes (21). A general increase in autophagy in C6R mice suggests that mHTT proteolysis disrupts the role of wt HTT in the regulation of autophagy and may thus be a major cause for autophagic dysfunction in HD.

## Results

### Hepatic HTT levels increase with age in YAC128 mice and can be reduced by acute fasting

While HD is primarily a disease of the CNS, the HTT protein is expressed ubiquitously and metabolic alterations, including changes in the liver, have been reported (22, 23). The liver heavily relies on autophagy to maintain its basal function (24), and dysfunction in autophagic pathways including autophagosomes devoid of cargo have been found in liver tissue from HD mouse models (5, 25). We therefore decided to assess HTT protein levels in the liver of the YAC128 HD mouse model and found a strong age-dependent increase in wt and mHTT protein that reached statistical significance at 12 months (Fig. 1A). On the other hand, C6R mice expressing mHTT resistant to caspase-6 cleavage at D586 showed no age-dependent alterations in wt or mHTT levels, suggesting that this change is specific to the expression of cleavable mHTT in YAC128 animals (Fig. 1A).

**Fig. 1:**
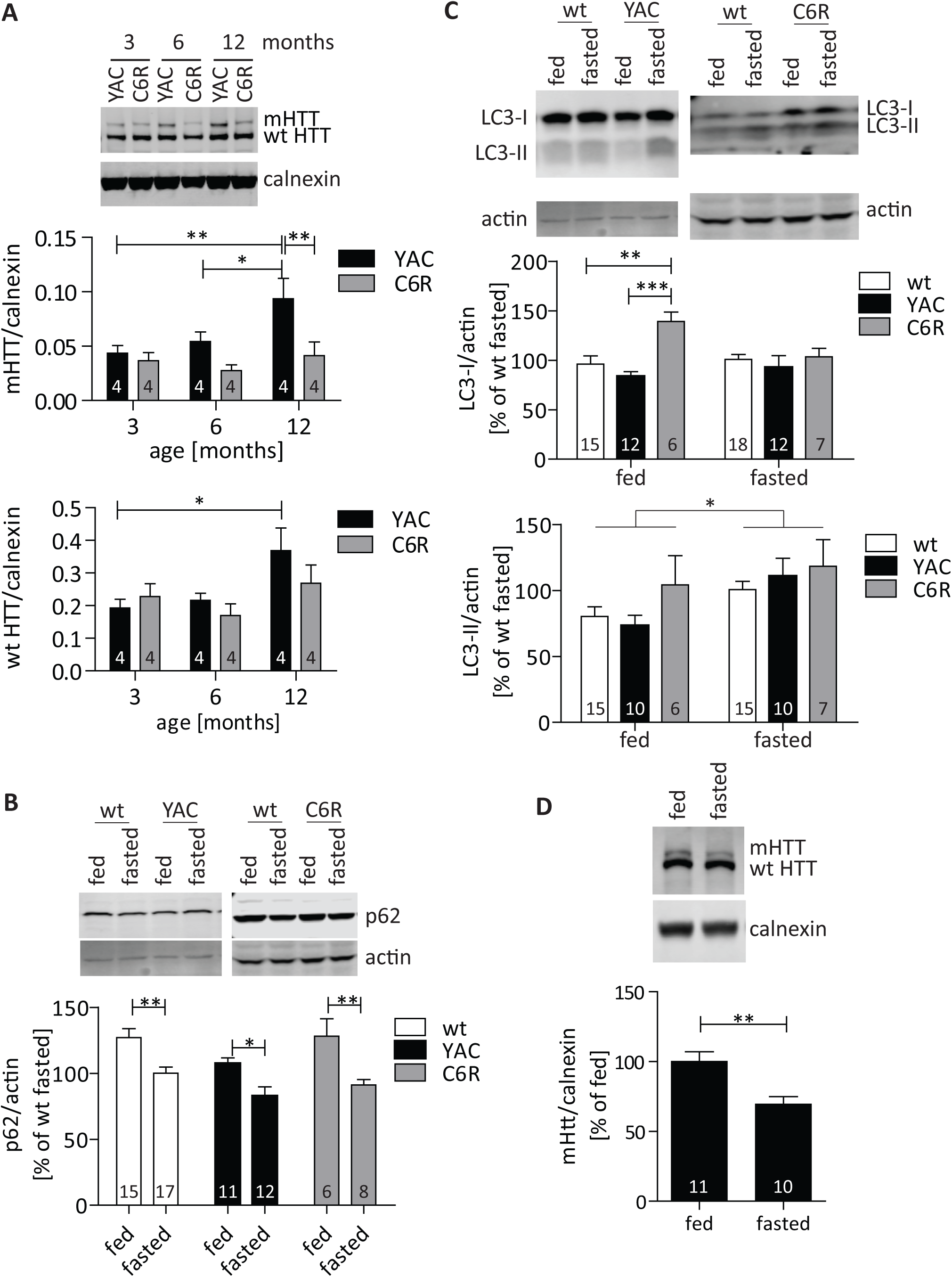
mHTT levels increase in aging YAC12S liver and can be reduced by fasting-induced autophagy. (A) Liver tissues from YAC128 and C6R mice at different ages were analyzed for HTT expression using the MAB2166 antibody. mHTT and wt HTT levels increase with age in YAC128, but not C6R liver tissue (mHTT ANOVA genotype p=0.0047, age p=0.0342; wt HTT ANOVA genotype p=0.3168, age p=0.0232). (B) - (D): 12 month old YAC128 and C6R mice, as well as their wt littermates, were subjected to a 24 h fasting period, sacrificed immediately and liver samples were compared to littermates with ad libitum access to food. p62 (B) and LC3 (C) protein levels were analyzed by Western blot. (B) Fasting reduces p62 in mice of all genotypes (ANOVA genotype p=0.0103, fasting p<0.0001). (C) C6R mice show elevated LC3-I levels at baseline, which normalize after fasting (ANOVA genotype p=0.0043, fasting p=0.3161). Fasting increases LC3-II levels in all animals (ANOVA genotype p=0.2012, fasting p=0.0151). (D) HTT protein levels in YAC128 liver were analysed by Western blot using the MAB2166 antibody. mHTT is significantly decreased after fasting. Representative blots and pooled quantification data with S.E.M. are shown, number of replicates are shown as insets. Statistical significance was determined by two-way ANOVA with Bonferroni’s post-hoc correction for (A) - (C), or Student’s t-test for (D). ^*^: p<0.05, ^**^: p<0.01, ^***^: p<0.001.

To study autophagic pathways *in vivo,* we next established a food deprivation paradigm. Food deprivation activates autophagy (26), and a fasting period of 24 h was sufficient to observe significant changes in hepatic levels of key autophagy proteins in wt as well as YAC128 mice: fasting decreased p62 levels, suggesting the induction of autophagy following food deprivation (Fig. 1B) (27). Furthermore, levels of the lipidated form of LC3, LC3-II, which decorates autophagosomes and is thus an indicator of autophagosome abundance (27), were specifically increased by fasting, while the non-lipidated form of the protein (LC3-I) was unaltered (Fig. 1C).

Interestingly, fasting-induced autophagy in the liver was paralleled by a significant reduction of mHTT protein in YAC128 mice (Fig. 1D), while the levels of wt HTT remained unchanged (Suppl. Fig. S1A). RNA levels of the mHTT transgene on the other hand were not affected by fasting, confirming that this intervention likely reduced mHTT protein through autophagic degradation (Suppl. Fig. S1B). Fasting also had an impact on hepatic mHTT protein levels in C6R mice, although the reduction was more subtle in this genotype (Suppl. Fig. S1C). This is not surprising, given the already low levels of mHTT protein in C6R compared to YAC128 mice. Nevertheless, the trend towards a further decrease suggests that fasting-induced autophagy can still lower mHTT even in C6R mice.

Metabolic dysfunction is a phenotype in HD mouse models overexpressing HTT, including the YAC128 model, and manifests as increased body weight gain with elevated levels of IGF-1 in the plasma (28). We found that fasting not only induced weight loss in both wt and YAC128 animals (Fig. 2A), but, more importantly, normalized the plasma levels of IGF-1 in YAC128 mice (Fig. 2B). Taken together, these findings demonstrate that fasting-induced autophagy mechanisms are intact and can be activated in the liver of YAC128 mice and suggest that the age-dependent accumulation of mHTT can be reversed by activating such protein degradation pathways simply through dietary changes.

**Fig. 2:**
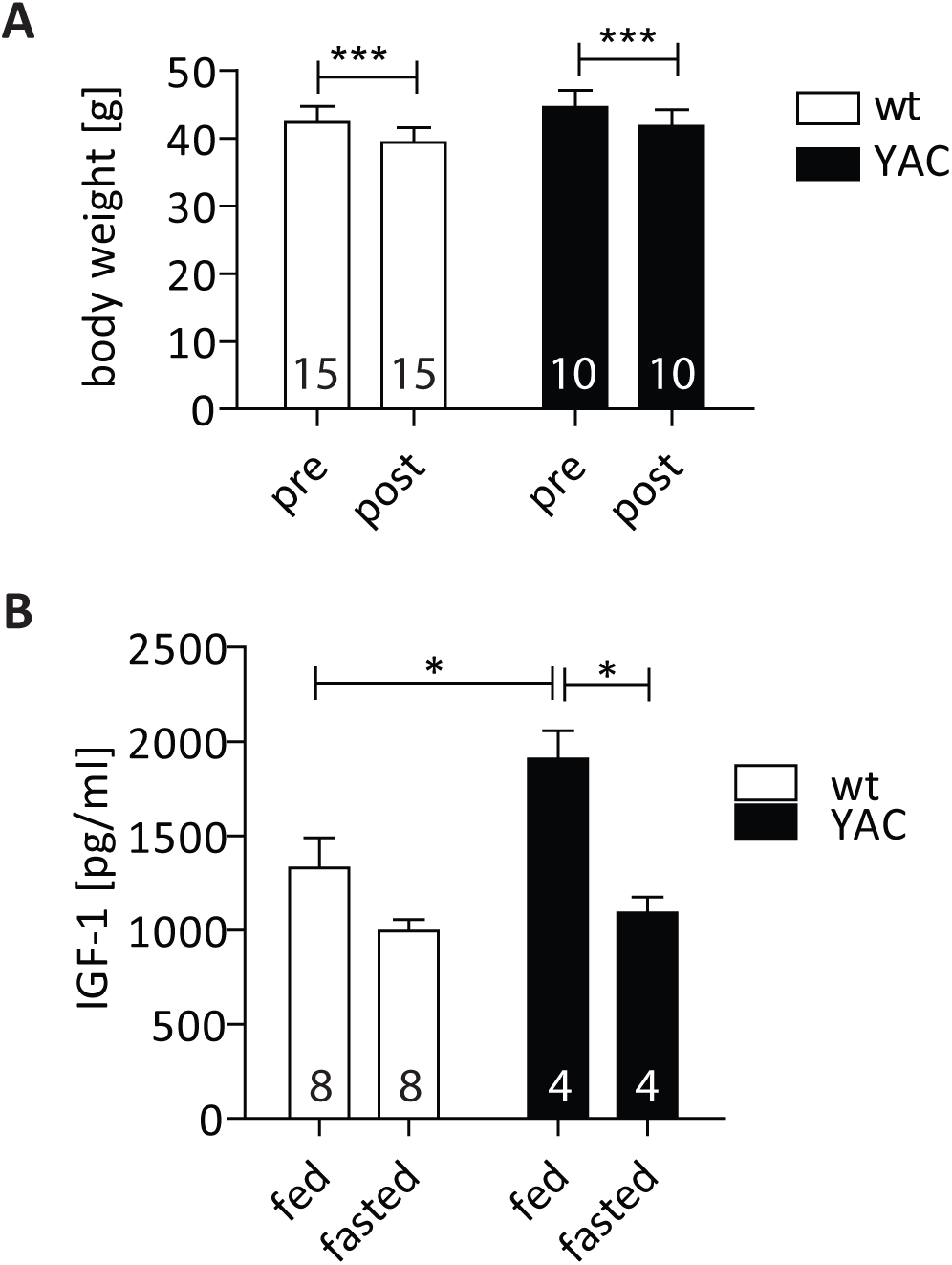
Fasting normalizes plasma IGF-1 levels in YAC128 mice. (A) Female YAC128 mice and their wt littermates were subjected to 24 h of fasting, and body weight was recorded pre- and post-fasting for each animal. No difference in weight loss was observed between genotypes (ANOVA genotype p=0.0968, fasting p< 0.0001). (B) IGF-1 levels in the plasma of fed and fasted YAC128 mice as well as their wt littermates were determined by ELISA. As reported previously, plasma IGF-1 levels are elevated in YAC128 mice at baseline (28). This phenotype is normalized after a 24 h fasting period (ANOVA genotype p=0.0241, fasting p=0.0004). Statistical significance in (A) and (B) was determined using two-way repeated measures ANOVA with post-hoc Bonferroni correction, number of replicates are shown as insets. ^*^: p<0.05, ^**^: p<0.01, ^***^: p<0.001.

### Prolonged regulation of food intake induces mHTT clearance in the brain

Recent studies have demonstrated that acute fasting also induces autophagy in the CNS (9, 10). We therefore examined LC3 and p62 levels in cortical tissues from fasted as well as mice fed ad libitum, and found that a 24h fasting period increased LC3-I, LC3-II and p62 protein levels in the cortex of both wt and YAC128 mice (Suppl. Fig. S2A). Interestingly, both the increased cortical p62 and LC3-I levels differ from our findings in the liver (Fig. 1B and C), which suggests different timing of autophagy induction in the two organs. Furthermore, unlike hepatic mHTT (Fig. 1D), cortical mHTT levels remained unaffected by a 24h fasting period (Supp. Fig. S2B). This is consistent with reports showing that autophagosome numbers first increase after 24h of fasting in the cortex, which is followed by their increased trafficking towards the neuronal soma and degradation, again reducing their steady-state level (9). However, in the liver different kinetics are expected, since autophagy induction happens earlier and autophagosomes do not have to travel long distances along neurites to be degraded. We therefore hypothesized that the lack of mHTT degradation after acute fasting may be due to the delayed induction of autophagy in the brain compared to the liver, and designed a scheduled feeding paradigm that allowed us to regulate the food intake of YAC128 mice for a longer period of time.

For scheduled feeding, the access to food was restricted to 6 h per day during the dark cycle, and mice were sacrificed after a week on scheduled feeding at the end of a fasting period. Long-term caloric restriction has beneficial effects on aging and neurodegenerative phenotypes (29, 30), and previous studies have implicated both mTOR inhibition and SIRT1 activation in these phenomena (29, 31-33). Consistent with these reports, we observed that scheduled feeding decreased phosphorylated mTOR (Fig. 3A), an indication of mTOR inhibition similar to the effect of nutrient deprivation (11). In addition, RNA levels of SIRT1 were increased in mice subjected to scheduled feeding (Fig. 3B), which can further contribute to the downregulation of mTOR signaling (33) and thus induce autophagy.

**Fig. 3:**
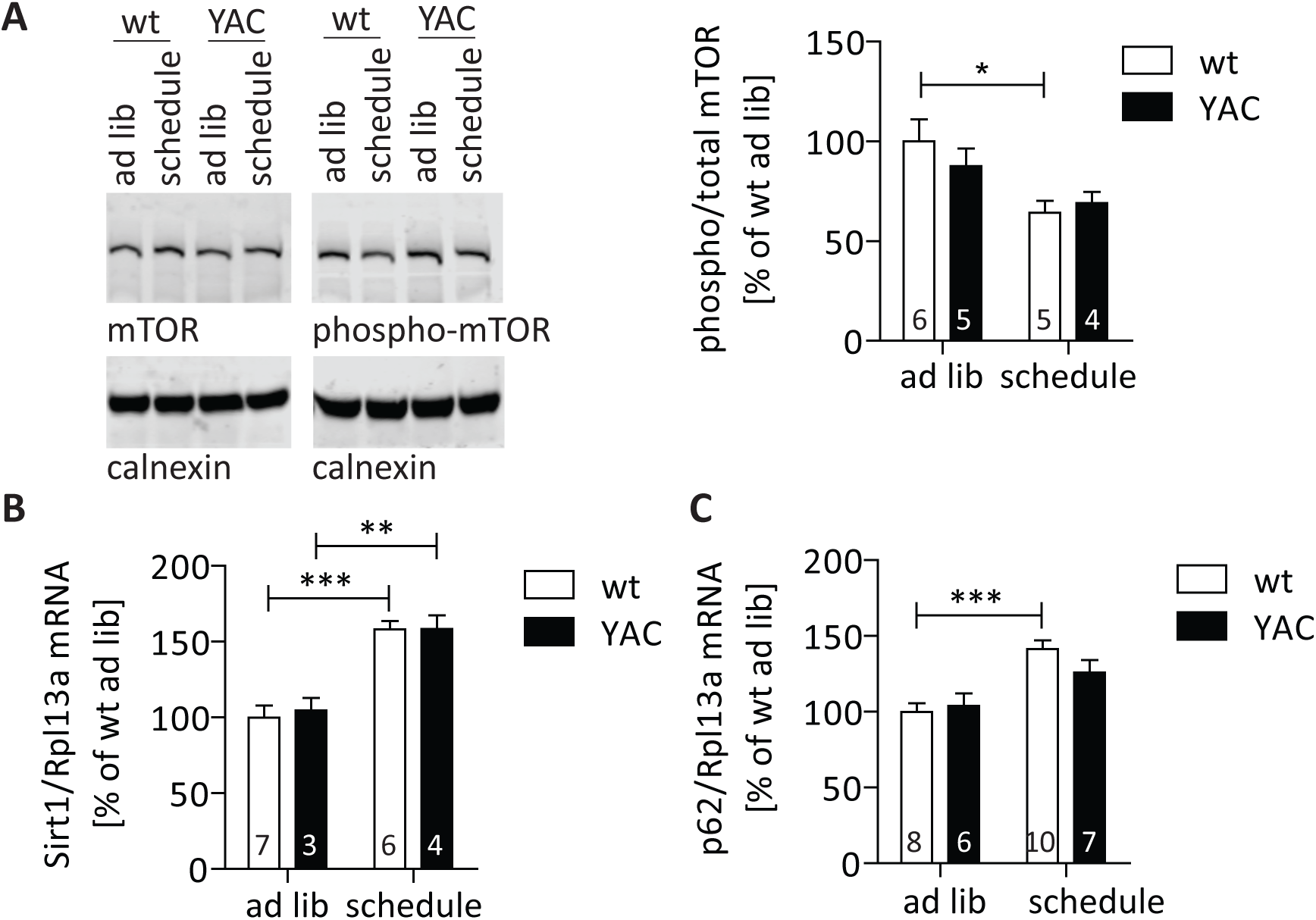
Scheduled feeding alters nutrient-sensing pathways in the brain. (A)-(C) YAC128 mice and their wt littermates were subjected to one week of scheduled feeding and compared to littermates with ad libitum access to food. (A) Scheduled feeding reduces the levels of phospho-mTOR, indicating an inhibition of mTOR signaling (ANOVA genotype p=0.6796, feeding p=0.0082). (B) Scheduled feeding increases the mRNA levels of SIRT1 in both wt and YAC128 mice, which may contribute to mTOR inhibition (33) (ANOVA genotype p=0.7651, feeding p<0.0001). (C) p62 expression is upregulated by scheduled feeding, a mechanism known to replenish p62 protein levels and maintain autophagic flux during long-term fasting (34, 35) (ANOVA genotype p=0.4121, feeding p<0.0001). Representative blots and pooled quantification data with S.E.M. are shown, number of replicates is shown as insets. Statistical significance was determined by two-way ANOVA with Bonferroni’s post-hoc correction. ^*^: p<0.05, ^**^: p<0.01, ^***^: p<0.001.

While the steady-state cortical LC3 or p62 protein levels were not altered by scheduled feeding (Suppl. Fig. S3A and B), we observed a significant induction of p62 expression at the RNA level (Fig. 3C). This is consistent with the previously demonstrated replenishing of p62 protein levels through increased expression during long-term fasting (34, 35). Furthermore, analysis of LC3 puncta by confocal microscopy demonstrated a significant decrease after scheduled feeding, consistent with increased turnover of autophagic vesicles decorated with LC3 (Fig. 4A). Interestingly, C6R mice exhibited a strong reduction in LC3 puncta even at baseline, which was not altered by scheduled feeding, suggesting that these animals already exhibit increased autophagic flux when fed a regular diet (Fig. 4A). Next, EM analysis was performed to detect the formation and loading of autophagosomes and autophagolysosomes in the cortex of scheduled-fed mice to assess induction of autophagy. In this experiment we observed predominantly autolysosomes with electron-dense content with some remaining ultrastructure (27), and only very few autophagosomes with a double membrane and visible content, and the quantification of these types of autophagic vesicles (AVs) has been proposed as a reliable method to assess autophagy induction (27). We found that the overall number of AVs per cell significantly increased after scheduled feeding, confirming that autophagosomes are successfully transported to the neuronal soma and cargo degradation increased (Fig. 4C). Consistent with this data we found that the scheduled feeding paradigm also significantly reduced the levels of cortical mHTT protein in YAC128 mice (Fig. 4C), similar to the effect of short-term fasting on mHTT the liver (Fig. 1D). No changes were observed for wt HTT protein (Suppl. Fig. S4A) or mHTT mRNA (Suppl. Fig. S4B). Taken together our data suggests that scheduled feeding increases mHTT clearance in the brain through the upregulation of autophagy.

**Fig. 4:**
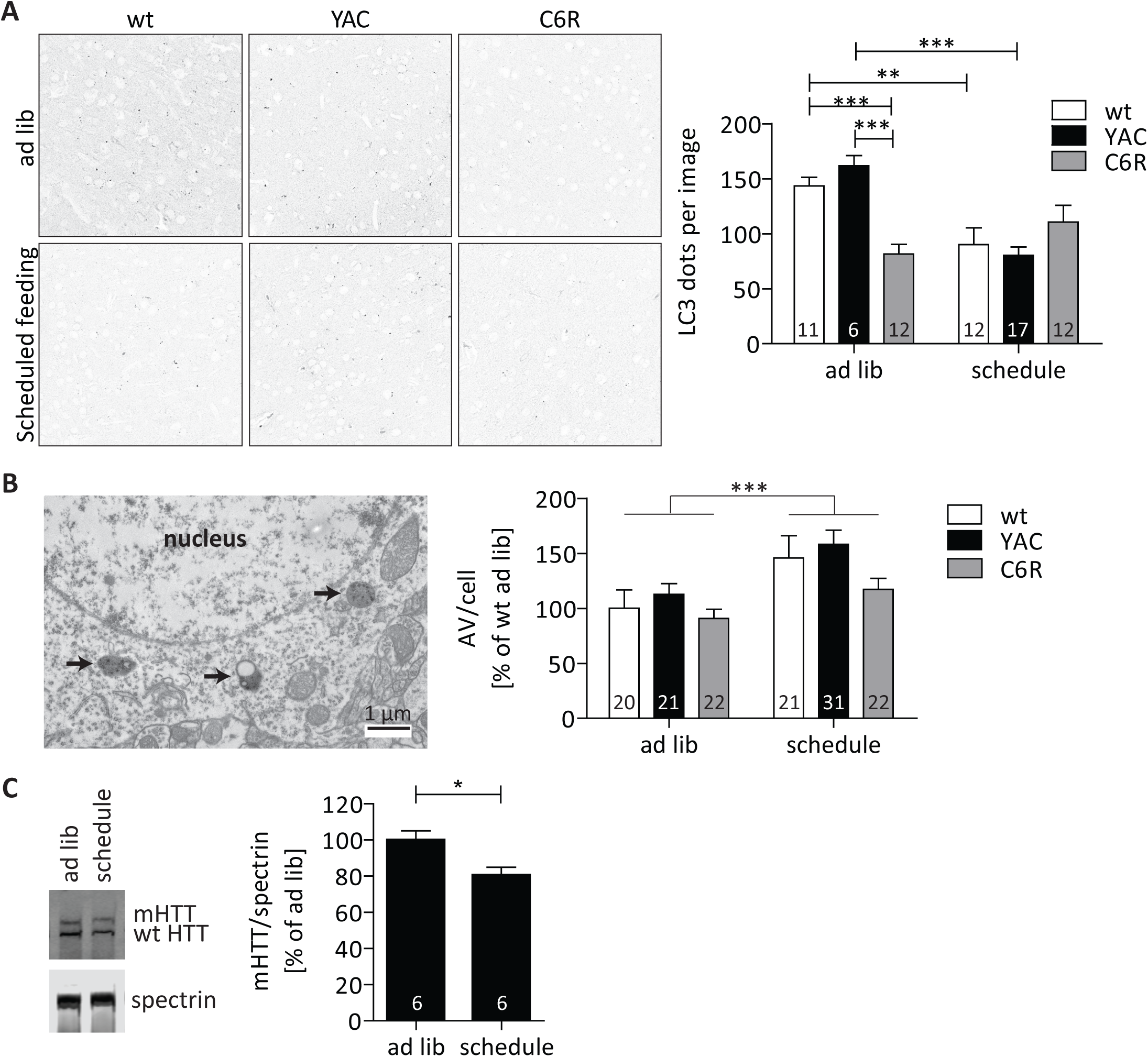
Scheduled feeding induces autophagy and lowers mHTT protein in the brain. (A) -(C) YAC128 and C6R mice as well as their wt littermates were subjected to one week of scheduled feeding and compared to littermates with ad libitum access to food. (A) Cortical brain sections were stained for LC3 and confocal images were analyzed. While mice of all genotypes showed reduced numbers of LC3 puncta after scheduled feeding, C6R mice already exhibited a significantly decreased LC3 puncta count at baseline (ANOVA genotype p=0.0815, feeding p=0.0006). (B) EM analysis of the motor cortex of the same YAC128, C6R and wt mice reveals an increase in autophagic vesicles (AV) surrounding the nuclei after scheduled feeding (ANOVA genotype p=0.0636, feeding p=0.0006). (C) Western blotting using antibody MAB2166 demonstrates that scheduled feeding reduces mHTT protein levels in the cortex of YAC128 mice. Representative images/blots and pooled quantification data with S.E.M. are shown, number of replicates is shown as insets. Statistical significance in (A) and (B) was determined by two-way ANOVA with Bonferroni’s post-hoc correction, in (C) by Student’s t-test. *: p<0.05, **: p<0.01, ***: p<0.001

### Preventing mHTT cleavage at D586 increases autophagy

While scheduled feeding lowered mHTT levels in YAC128 mice, increased autophagic clearance through forced circadian feeding does not sufficiently explain our finding that wt and mHTT do not accumulate with age in C6R liver (Fig. 1A). C6R mice exhibit the same age-dependent weight gain as YAC128 animals (Suppl. Fig. S5), suggesting similar metabolic phenotypes in both mouse models (23, 28). However, it has been suggested that the autophagy-promoting function of HTT itself may be lost in HD due to proteolytic events separating the N- from the C-terminal region of the protein (36, 37). In C6R mice, the analysis of hepatic autophagy proteins revealed elevated levels of LC3-I at baseline, which are reduced upon fasting for 24 h (Fig. 1C). This suggests that a larger pool of LC3-I is available, which can be immediately converted into LC3-II and promote autophagy upon a change of nutritional status. Furthermore, we found that the steady-state levels of LC3 puncta in the cortex were significantly lower in C6R compared to both wt and YAC128 mice (Fig. 4A), suggesting altered autophagy flux in the brain.

We therefore assessed autophagy proteins in primary neuronal cultures using bafilomycin, an inhibitor of autophagic flux, which allows for a more accurate quantification of LC3-II in the absence of its turnover (27). Consistent with results from liver tissues, LC3-I levels were also dramatically increased in neurons expressing C6R mHTT, while LC3-II levels were similar to wt neurons (Fig. 5A). In addition, baseline levels of p62 were elevated in C6R neurons, but both LC3-II and p62 further increased after bafilomycin treatment (Fig. 5A), suggesting that autophagic flux was not blocked, but rather increased overall (27). Conversely, the data for YAC128 neurons showed a decrease in autophagic flux, since p62 levels were significantly lower than in wt and LC3-II showed a trend towards a reduction after bafilomycin treatment (Fig. 5A). These findings are consistent with the hypothesis that cleavable mHTT impairs autophagosome formation, whereas the expression of cleavage-resistant C6R mHTT not only repairs, but boosts autophagy (36).

**Fig. 5:**
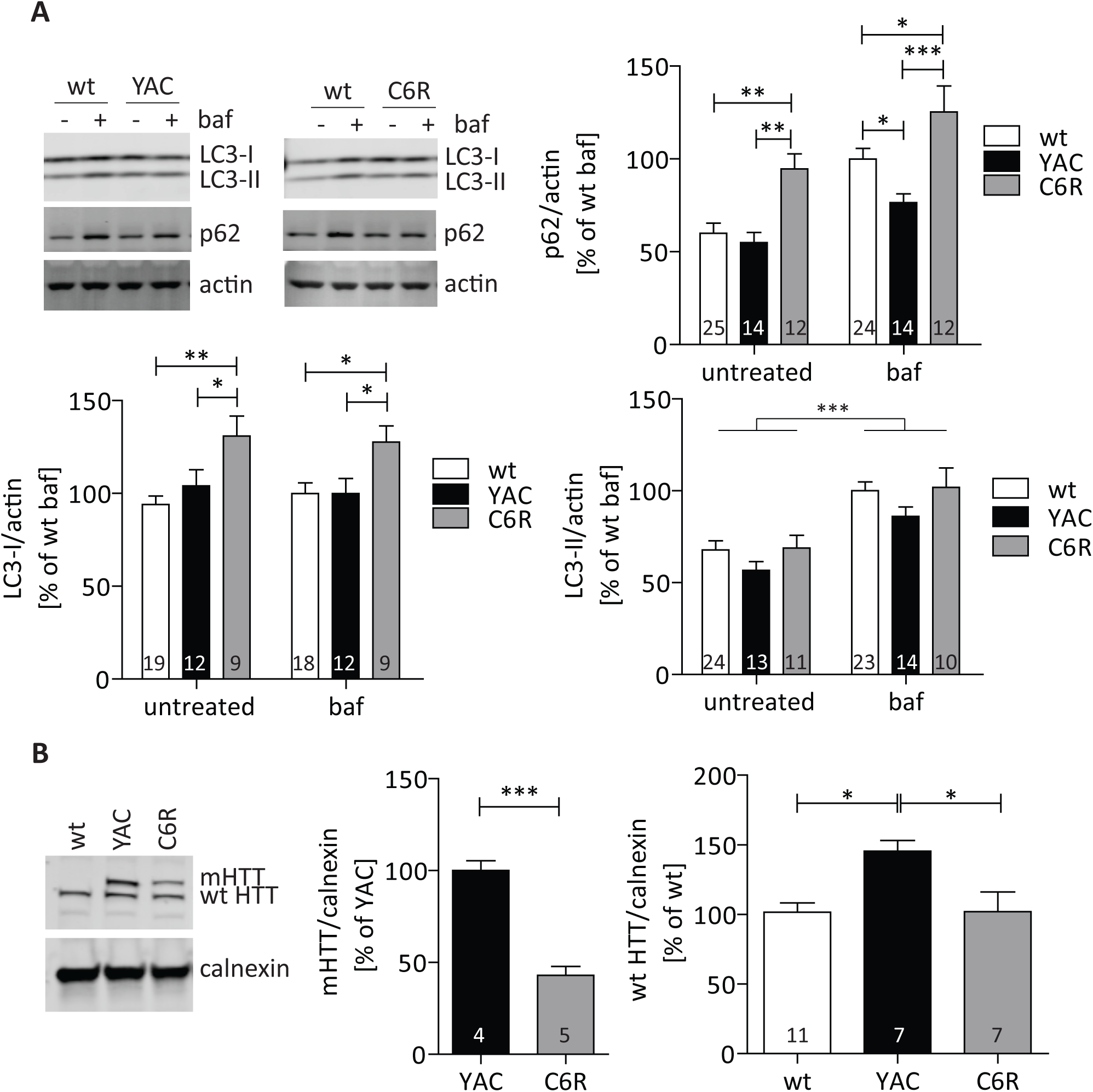
Preventing mHTT cleavage restores its autophagy-promoting function. (A) Primary neuronal cultures from YAC128, C6R or wt littermate embryos were treated with 10 nM bafilomycin for 2 h or DMSO as control. Levels of p62, LC3-I and LC3-II were analyzed by Western blot. Both p62 and LC3-II levels increase after bafilomycin treatment in cultures of all genotypes, demonstrating that autophagic flux is intact. C6R neurons show increased baseline levels of p62 and LC3-I, consistent with the increased LC3-I levels observed in the livers of C6R animals (Fig. 1C). p62: ANOVA genotype p<0.0001, bafilomycin p<0.0001, LC3-I: ANOVA genotype p=0.0002, bafilomycin p=0.9352, LC3-II: ANOVA genotype p=0.0562, bafilomycin p<0.0001. (B) Primary neuronal cultures derived from YAC128, C6R or wt littermate embryos were analyzed for HTT protein levels with the MAB2166 antibody. YAC128 neurons exhibit higher levels of mHTT than C6R cells and also show increased wt HTT protein compared to both wt or C6R neurons (wt HTT ANOVA p=0.0088). Representative blots and pooled quantification data with S.E.M. are shown, 3-5 independent cultures were analyzed, number of replicates are shown as insets. Statistical significance was determined using two-way ANOVA with post-hoc Bonferroni correction for (A), one-way ANOVA with post-hoc Tukey correction for (B): wt HTT, or Student’s t-test for (B): mHTT. *: p<0.05, **: p<0.01, ***: p<0.001.

Next, we sought to determine whether these changes in autophagy alter HTT levels. Surprisingly, we found that YAC128 neurons contained twice as much mHTT and 50% more wt HTT compared to C6R or wt neurons (Fig. 5B). Both protein and mRNA levels of mHTT in C6R and YAC128 mice were previously found to be comparable in the CNS (21, 38). However, a lower incidence of mHTT inclusions in C6R compared to YAC128 animals by immunohistochemistry has been shown previously (21, 39), which may reflect differences in mHTT clearance by autophagy.

### The C6R mutation rescues the interaction of mHTT with the autophagy receptor p62

To further investigate the role of HTT as autophagic cargo, we next assessed its interaction with the autophagy receptor p62. This protein is links both wt and mHTT to LC3-II and the autophagosome, thus mediating their autophagic degradation (25, 40). Indeed, our co-immunoprecipitation experiments showed that the presence of the expanded polyglutamine tract significantly reduced the binding efficiency of HTT to p62 (Fig. 6A). Co-transfecting p62 and cleavable, C6R or a 586aa fragment of mHTT revealed that C6R mHTT interacts approximately twice as strongly with p62 as cleavable mHTT (Fig. 6B), which was confirmed with co-immunoprecipitation of HTT with p62 (Suppl. Fig. S6). Interestingly, the interaction between p62 and a mHTT-586aa fragment was barely detectable, suggesting that the major p62 interaction site in HTT is located C-terminal of the D586 caspase cleavage site (Fig. 6B), as previously described (37), or multiple interaction domains are required that are separated by proteolysis.

**Fig. 6:**
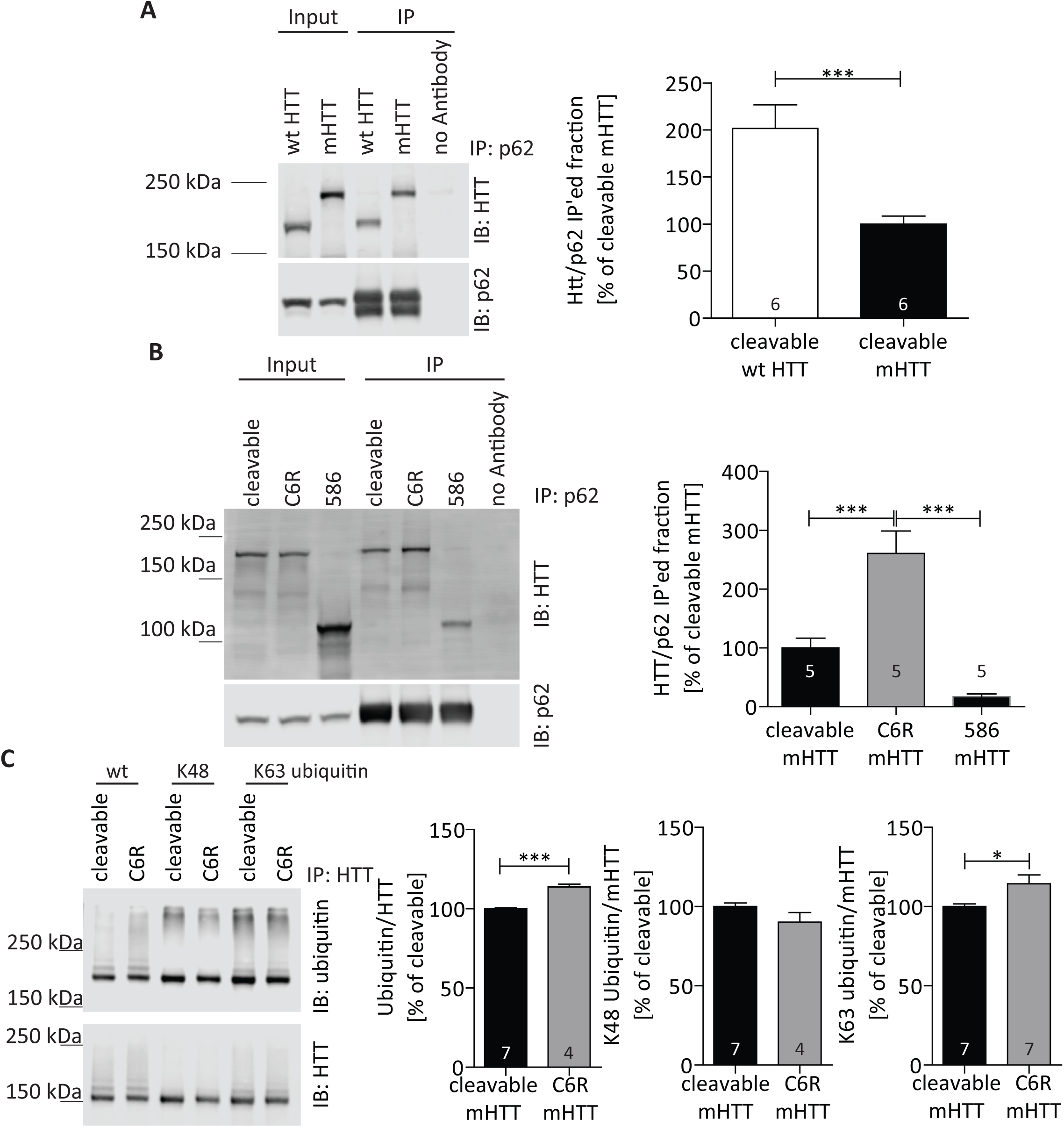
The C6R mutation improves the binding of mHTT to the autophagy adapter p62. (A) and (B) COS-7 cells were cotransfected with HTT aa1-1212 (wt or mutant, cleavable or C6R) or mHTT aa 1-586 and p62 as indicated. After immunoprecipitation of p62, the ratio of co-immunoprecipitated HTT was quantified (normalized to input to control for transfection efficiency). (A) Cleavable mHTT exhibits reduced binding to p62 compared to the wt HTT construct. (B) The binding of C6R mHTT to p62 is increased compared to cleavable mHTT, whereas the mHTT construct truncated at aa586 binds very poorly to the autophagy adapter protein (ANOVA p<0.0001). (C) COS-7 cells were cotransfected with mHTT aa1-1212 (cleavable or C6R) and HA-tagged wt, K63 or K48 ubiquitin (allowing all, only K63 or only K48 linkage to target proteins) as indicated. After immunoprecipitation of HTT, the ratio of co-immunoprecipitated ubiquitin/HTT was quantified (normalized to input to control for transfection efficiency). C6R mHTT preferentially bound to K63 ubiquitin, and overall ubiquitin binding was also increased for this construct. Blots and quantification data with S.E.M. from a representative of 3 independent experiments are shown, number of replicates is shown as insets. Statistical significance was determined by Student’s t-test for (A) and (C) or one-way ANOVA with Tukey’s post-hoc correction for (B). ^*^: p<0.05, ^**^: p<0.01, ^***^: p<0.001.

p62 interacts with ubiquitinated substrates, and preferentially those that are linked to ubiquitin through lysine 63 (K63) (41, 42). Co-transfection of cleavable or C6R mHTT with either wt ubiquitin, or ubiquitin mutants that can only bind their target proteins through lysine 48 (K48 ubiquitin) or lysine 63 (K63 ubiquitin), revealed that C6R mHTT co-immunoprecipitated with significantly more wt ubiquitin (wt ubiquitin, Fig. 6C). Interestingly, the interaction with K48 ubiquitin was equal between cleavable and C6R mHTT, but K63 ubiquitin preferentially co-immunoprecipitated with C6R mHTT, indicating that the K63 linkage is preferred in the presence of the C6R mutation (Fig. 6C). Increased K63-ubiquitination of C6R mHTT would thus be expected to mediate increased p62 binding and may therefore account for its preferential autophagic clearance.

## Discussion

The expansion of the CAG tract in HTT is the single cause for HD, and recent efforts in therapeutic development have therefore focused on different strategies to lower the levels of mHTT (3). Here we show that mHTT clearance can be enhanced in an HD mouse model by scheduled feeding as well as through reduced proteolysis. These findings may have significant implications for clinical trials using mHTT lowering strategies that are currently under way: firstly, baseline levels of mHTT clearance may differ between HD patients due to dietary and other environmental factors influencing autophagy, and secondly, clearance may be decreased in advanced disease stages or with advanced age of the patient, as the activity of proteolytic enzymes increases with aging (43, 44). Indeed, it was recently demonstrated that mHTT levels increase with disease progression in patient CSF (45).

Here we report that wt and mHTT accumulates with age in the liver of YAC128 mice, which may be caused by decreasing autophagic capacity (25). Hepatic mHTT expression is associated with decreased function of the urea cycle, lowered expression of PGC-1α and metabolic dysfunction (46-48). In addition, aggregates of mHTT have been found in the livers of the R6/2 HD mouse model (49). Interestingly, we do not observe an age-dependent accumulation of mHTT in the liver of C6R mice. The C6R mice are remarkable insofar as they express mHTT, but do not exhibit any HD-like phenotypes (21, 43). Our results suggest that this phenomenon may be due to increased autophagy and autophagic degradation of mHTT, leading to significantly lower steady-state levels of mHTT protein. While we show that C6R mHTT is a better substrate for p62-mediated autophagy, the restoration of autophagic flux likely plays an additional role, since C6R liver tissue shows high levels of LC3-I, which is rapidly turned over upon fasting. Supporting this hypothesis, increased autophagy and reduced mHTT levels have been demonstrated previously in HD mouse models crossed to a C6^-/-^ line, an intervention that reduces (but does not completely abolish) mHTT cleavage at D586 (50, 51).

Recent studies have shown that in addition to being a substrate for autophagic clearance, HTT can also promote autophagy. The C-terminus of the protein can act as a scaffold similar to Atg11, and promote autophagosome formation (37). This novel function of HTT is supported by additional data demonstrating that myristoylation of a C-terminal HTT fragment at G553 leads to a dramatic increase of autophagosome formation *in vitro* (52). While cleavage at D552 opens G553 up for myristoylation and may therefore be required for HTT-promoted autophagy, additional proteolysis at D586 would shorten the C-terminal fragment and may thus interrupt HTT’s scaffold function (Fig. 7). A recent study has indeed demonstrated that HTT N-terminal fragments interact with the HTT C-terminus after a single cleavage event, but this interaction is disrupted in the case of multiple proteolytic events (53). We therefore propose a model in which the negative effects of mHTT N-terminal fragments on autophagosome formation, transport and fusion are repaired in C6R mice in which the second mHTT cleavage at D586 is prevented (Fig. 7). This furthermore boosts the normal function of the intact HTT C-terminus in promoting autophagosome formation and increased binding to p62 allows for more efficient degradation of mHTT itself (Fig. 7).

**Fig. 7:**
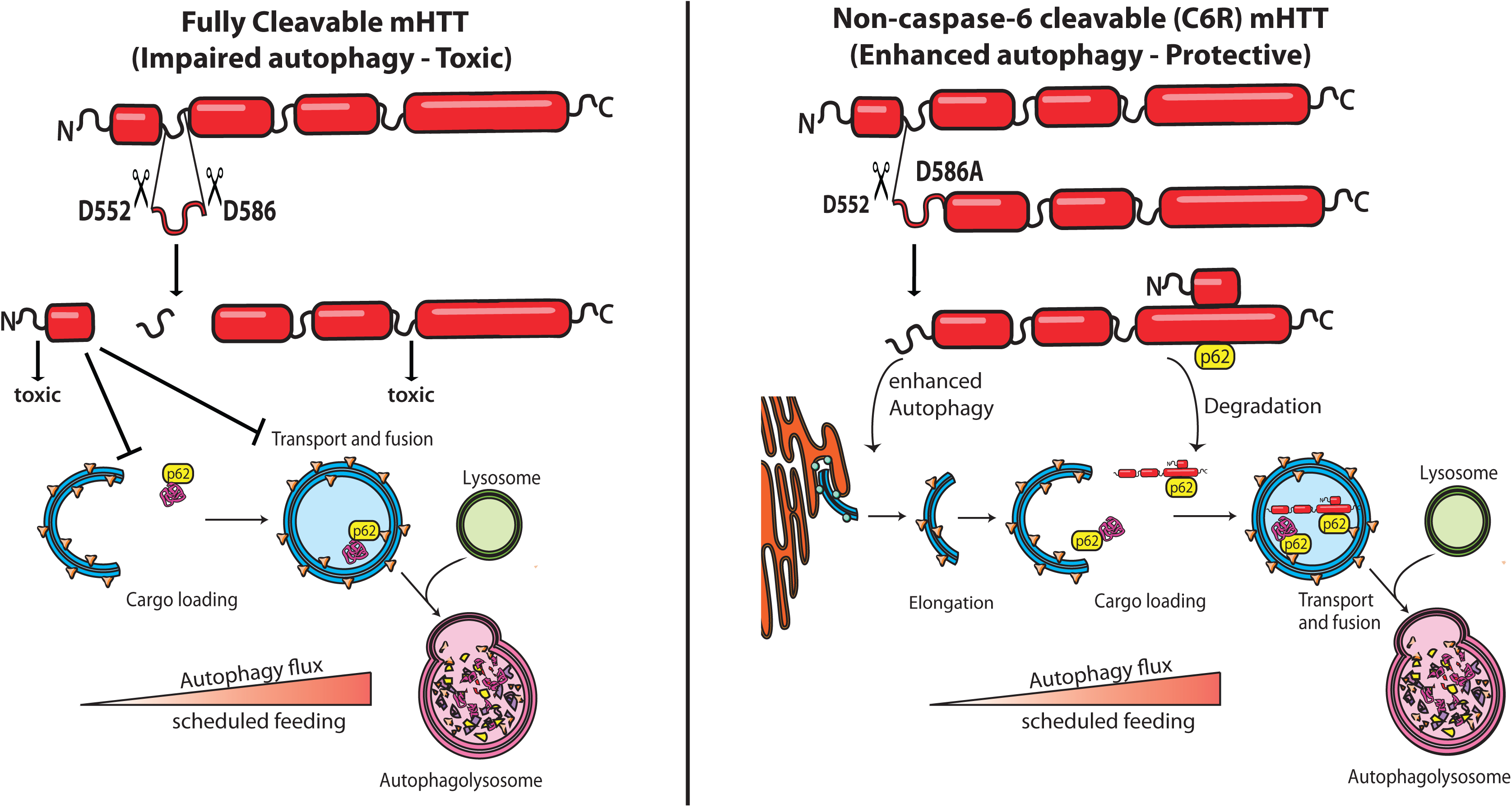
Schematic representation of the effects of cleavable or C6R mHTT on autophagy. mHTT is subject to proteolysis by different proteases, with a number of cleavage sites clustering in the PEST2 domain (64). Cleavage at D552 and D586 liberate a small fragment that is myristoylated at G553 (52) and induces autophagosome formation. However, both the N-terminal and C-terminal fragments resulting from mHTT double cleavage are toxic and interfere with autophagic cargo loading and autophagosome transport and fusion (25, 37, 53, 65). In the C6R mice, cleavage at D586 is prevented, and we observed enhanced autophagy as well as improved degradation of C6R mHTT in cells and tissues from C6R mice. In both YAC128 and C6R mice, autophagic flux and mHTT degradation can be enhanced in peripheral tissues by fasting and in the CNS by scheduled feeding.

mHTT aggregates accumulate with aging, and HD is a progressive disease that in the majority of patients only manifests in adulthood. While caloric restriction can slow aging in a large variety of animal models (54) and upregulate key transcription factors such as *SIRT1* that are beneficial in different neurodegenerative conditions including HD (29, 31, 32), such an intervention is not advisable for human patients, since HD already leads to a significant reduction in body weight (55) which may be exacerbated by further calorie reduction. However, we show here that scheduled feeding is sufficient to upregulate *SIRT1* expression and activate the mTOR pathway in a mouse model of HD. Circadian rhythms are disrupted in HD patients as well as in animal models of the disease, and this phenotype can be ameliorated by regulating food intake (12). Interestingly, circadian patterns are improved by this intervention even in mice at late stages of the disease (12). This is in line with our findings that fasting induces mHTT clearance in the liver in 12 month old YAC128 mice, which have already accumulated significant amounts of the mutant protein. Since autophagy follows a circadian pattern in the brain (9), it is possible that the disruption of circadian rhythms in neurodegenerative disease may cause autophagic dysfunction and contribute to the accumulation of autophagy substrates such as mHTT. Furthermore, there is evidence that treating disruptions in circadian rhythm may ameliorate symptoms such as depression, anxiety and cognitive dysfunction in human HD patients (13), and our data suggest that such an intervention additionally has the potential to lower mHTT protein levels through increased autophagy.

## Materials and Methods

### Statistics and animal models

All mouse experiments were carried out in accordance with protocols (Animal protocol A07-0106) approved by the UBC Committee on Animal Care and the Canadian Council on Animal Care.

YAC128 (line HD53 (56)) and C6R (line W13 (21)) mice are on a FVB/N background, mixed sexes were analyzed. Cortex and liver tissue was dissected and snap-frozen on dry ice for protein analyses. Sample sizes were chosen based on extensive experience with biochemical differences between YAC128 mice and their WT littermates for experiments using mouse tissues (21, 23, 28, 38, 43, 51, 57). Cell culture experiments were repeated independently at least three times to ensure reproducibility. Samples were only excluded if technical issues were apparent (i.e. bubble on a Western blot) or if determined statistical outliers using Grubb’s outlier test (α = 0.05, no more than one sample per group was excluded).

For randomization, mice were assigned numbers not related to genotype. Scientists performing experiments were blinded for genotypes, unless it was necessary to ensure the appropriate order of samples on a gel. Data analyses was performed by a separate person in possession of the genotype information. For image analysis of electron microscopy and confocal microscopy data, unblinding was performed after all quantitation was complete.

Statistical significance was assessed using Student’s t-test for comparison of two groups, one-way ANOVA with post-hoc Tukey’s correction for the comparison of one variable between more than two groups, and two-way ANOVA with post-hoc Bonferroni correction for the comparison of two variables between groups. Variances between groups were similar. All analyses were performed using the GraphPad Prism 5.01 software package.

### Generation and treatment of primary neuronal cultures

YAC128 (line 53) and C6R (line W13) embryos, as well as their wt littermates, were collected on day 15.5-16.5 of gestation. Brains were extracted and transferred to Hibernate E (Invitrogen) for up to 24h, during which time samples from the remaining embryonic tissues were genotyped (58).

Cortices were micro-dissected in ice-cold Hank’s balanced salt solution (HBSS+; Gibco), then diced and pooled for each genotype. Cells were dissociated with 0.05% trypsin-EDTA (Gibco), followed by neutralization with 10% fetal calf serum in neuro basal medium (NBM+) and DNAse I treatment (153U/mL). Tissue was triturated with a pipette five to six times. Cells were plated on poly-D-lysine coated 6-well plates with 2 ml of Neurobasal media (Gibco #21103-049), B27 (Gibco #17504-044), 100 U/mLpenicillin-streptomycin (PS) (Gibco), 0.5 mM L-glutamine and maintained at 37°C, 5% C02 with humidity. Cells were fed with 200 mL fresh medium every fifth day. On day 9-11 in culture, cells were treated with 10 nM bafilomycin A1 (Cayman Chemicals) or DMSO for 2 h. Cells were harvested by scraping and lysed for Western blot analysis as described (59).

### Western blotting

Western blots were performed on samples lysed in SDP lysis buffer (50 mM Tris pH 8, 150 mM NaCl, 1% Igepal with ‘Complete’ protease inhibitor cocktail (Roche)). Protein concentration was measured using the DC protein assay kit (Bio Rad, USA) and equal amounts were separated on 7% Bis-Tris gels for the detection of HTT, or 4-12% gradient gels (Invitrogen, USA) for the detection of LC3 and p62. Protein was transferred to PVDF Immobilon-FL membranes by electroblotting and membranes were developed with primary antibodies in 5% bovine serum albumin/phosphate buffered saline. The following antibodies were used for immunoblotting: anti-HTT BKP1 (1:100), anti-HTT 2166 (1:1000) from Millipore (MAB2166), anti-p62 (1:1000) from ENZO (BML-PW9860), anti-LC3b (1:1000) from Cell Signaling Technologies (2775), anti-mTOR from Cell Signaling Technologies (2983), anti-phospho-mTOR from Cell Signaling Technologies (5536), anti-HA (1:1000) from COVANCE (MMS-101R), anti-Actin (1:10 000) from Sigma (A2103), anti-calnexin (1:5000) from Sigma (C4731). Fluorescently labelled secondary antibodies conjugated to either 700 or 800 IR dye (1:5000; Rockland, USA) and the LiCor Odyssey Infrared Imaging system were used for detection.

### Filter test

The detection of insoluble mHTT aggregates in mouse brain lysates with the filter test method was performed as previously described (57).

### FRET

Cortical lysates were prepared with 1 mL/100 mg tissue of FRET lysis buffer (10 mM HEPES pH 7.9, 1.5 mM MgCl_2_, 10 mM KCl, 420 μM Pefabloc, 100 μM DTT, and 1x ‘Complete’ protease inhibitor cocktail (Roche)) with an automated tissue homogenizer for 30 sec and then an additional 20 strokes with a Dounce homogenizer. Lysates were incubated for 1 h on ice, centrifuged at 2000 x *g* for 15 min at 4°C, and the supernatant was used for the detection of soluble mHTT levels by FRET as described previously (38).

### qRT-PCR

RNA was extracted using the PureLink mini RNA extraction kit (Life Technologies). RNA was treated with DNase I (Invitrogen) and 500 ng of RNA were reverse transcribed using Superscript III (Invitrogen) and oligo-dT primers according to manufacturer’s instructions to generate cDNA for qRT-PCR. The PCR was run with SYBR Green Power master mix (Applied Biosystems) on the ABI Prism 7500 Sequence Detection System.

Each sample was run in triplicate. Relative gene expression was determined by using the ΔΔC_T_ method, normalizing to *Rpl13a* mRNA levels. The following primers were used:

Human *Htt* forward: 5’-GAAAGTCAGTCCGGGTAGAAC-3’

Human *Htt* reverse: 5’-CAGATACCCGCTCCATAGCAA-3’

mouse *Rpl13a* forward: 5’-GGAGGAGAAACGGAAGGAAAAG-3’

mouse *Rpl13a* reverse: 5’-CCGTAACCTCAAGATCTGCTTCTT-3’

mouse *Sirt1* forward: 5’-CAGTGTCATGGTTCCTTTGC-3’

mouse *Sirt1* reverse: 5’-CACCGAGGAACTACCTGAT-3’

mouse *p62* forward: 5’-CTCAGCCCTCTAGGCATTG-3’

mouse *p62* reverse: 5’-TCCTTCCTGTGAGGGGTCT-3’

### COS-7 cell transfection and immunoprecipitation

COS-7 cells were maintained in DMEM with 10% FBS, 1% L-glutamine, 100 U/mL penicillin and 0.1 mg/mL streptomycin at 37°C and 5% C02 in a humidified PMID 20495340 incubator. Cells were transiently cotransfected with the aa1-1212 15Q, aa1-1212 128Q (cleavable), aa1-1212 138Q-C6R or aa1-586 Q128 HTT constructs (60) together with RFP-p62 (obtained from Addgene (61)) or HA-ubiquitin wt, K63 or K48 (obtained from Addgene (62)). The Xtreme gene 9 transfection reagent (Roche Applied Science, Quebec, Canada) was used according to the manufacturer’s protocol.

The day after transfection, cells were treated with 100 nM bafilomycin for 4 h to prevent HTT-p62 and HTT-ubiquitin complexes from degradation. Cell lysates were prepared in SDP buffer and immunoprecipitated over night at 4°C using anti-p62 (MBL PM045) or anti-HTT 2166 antibodies (Millipore MAB2166). Immunoprecipitates and cell lysates were subjected to SDS-PAGE and Western blot as described above.

### Transmission electron microscopy (TEM)

Mice were anesthetized with avertin and injected with 15 μL of heparin intracardially. Mice were perfused with 4% paraformaldehyde and 0.125% glutaraldehyde for 20 min at a rate of 6 mL/min. Brains were dissected and left overnight in fixative at room temperature. 400 μm sections were cut on a vibratome and 1 mm^2^ tissue blocks of motor cortex were dissected. Postfixing, embedding, sectioning and staining were performed at the University of British Columbia Bioimaging facility. Briefly, samples were rinsed in 0.1 M sodium cacodylate buffer and secondary fixed in 1% osmium tetroxide with 1.5% potassium ferricyanide in 0.1 M sodium cacodylate for 2 h. Tissue was washed two times in distilled water and dehydrated in a series of ethanol dilutions, followed by graded resin infiltration and embedding. Ultrathin sections were prepared on a Leica Ultracut 7 using 45 degree diamond histoknife. Thin sections were counterstained with Sato’s lead. Images were taken using a Hitachi H7600 Transmission Electron Microscope and analyzed using Image J. 10-15 cells per mouse were identified and pictures taken systematically around the nucleus, covering all visible cytoplasm. 4-5 mice per genotype from 2-4 separate litters of YAC and wt mice were analyzed in a randomized and blinded fashion. AV were identified as either autophagosomes with a double membrane and visible cytoplasmic content such as mitochondria, or autolysosomes (vesicles with electron-dense content and some remaining ultrastructure) (27) and counted manually. The number of AV/cell was calculated and normalized to wt littermates. Imaging and counting were performed by separate blinded investigators.

### Confocal microscopy and image analysis

Slides were imaged on a Leica SP5 laser-scanning confocal microscope with a 63X immersion plan-apochromat objective. Fixed and stained mouse cortices were imaged at 100 Hz with a 1024×1024 pixel scan format, a zoom factor of 1 and a pinhole size of 75 μm. LC3 staining was imaged using the 543 laser at 15% laser power (50% intensity and 100% gain) and DAPI was imaged using the UV laser at full laser power (25% intensity and 10% gain).

Background subtraction on the LC3 image stack was performed using the rolling ball method using a radius of 10 pixels. A threshold mask was then applied to the image stack based on intensities ranging from 100-255 to generate binary images. Subsequently, ImageJ (63) automated particle analysis was performed on the image stack and particle counts, size and area were measured for all images.

### 24 h fasting

Litters were split into even numbers of mice remaining on ad libitum diet and mice undergoing the fasting protocol. Mice assigned to the fasting cohort were weighed immediately before the start of the procedure and moved to clean cages. Food was withdrawn starting at the beginning of the light cycle, the animals had free access to drinking water at all times. At 15 h, 18 h and 21 h after the start of food deprivation animals were monitored for signs of dehydration and body weight loss and were injected with 0.5 mL sterile, warm saline solution subcutaneously if deemed necessary. After a 24 h fasting period, the mice were sacrificed immediately by lethal avertin injection and tissues were removed for analysis as described above.

### Scheduled feeding

Litters were divided into even numbers of mice remaining on ad libitum diet and mice undergoing the fasting protocol. Mice assigned to the scheduled feeding cohort were weighed immediately before the start of the procedure and moved to clean cages. For a total period of one week, access to food was restricted to the first 6 h of each dark cycle. The animals had free access to drinking water at all times. At the end of the last fasting period, the mice were sacrificed immediately by lethal avertin injection and tissues were removed for analysis as described above.

### IGF-1 and body weight measurements

IGF-1 levels in plasma were determined by ELISA as described previously (28). For body weight assessments, female mice were weighed in the housing facility at different ages using a digital scale.

## Author contributions

DEE designed and performed experiments, coordinated author contributions and wrote the manuscript. DDOM designed and performed experiments, provided intellectual input and edited the manuscript. XQ, SL, NSC, NHS, YTNN, KV, SE and SF performed experiments. MRH supervised the project and edited the manuscript.

## Acknowledgements

This work was supported by the Canadian Institutes of Health Research (CIHR 20R90174) and a sponsored research agreement with Teva Pharmaceuticals. The authors thank Dr. Amber Southwell for establishing the scheduled feeding paradigm, Dr. Ana-Maria Cuervo for assistance in the identification of AVs and Mandi Schmidt for assistance with image analysis. We thank Piers Ruddle, Mahsa Amirabbasi, Sheng Yu, Mark Wang, Yun Ko, Yuanyun Xie and Qingwen Xia for their technical support. DDOM and NHS were supported by postdoctoral fellowships from CIHR. DDOM was also supported by Michael Smith Foundation for Health Research and the Bluma Tischler Fellowship from UBC. SL was supported by a doctoral scholarship from CIHR. NSC was supported by postdoctoral fellowships from CIHR and the James Family.

## Competing financial interests

M.R.H. is an employee of Teva Pharmaceuticals, Inc. Teva did not play a role in the design, analysis or interpretation of this study. All other authors declare no competing financial interests.

